## Supplementary File 1 for "In situ mutational screening and CRISPR interference define *apterous* cis-regulatory inputs during compartment boundary formation"

OR463 sequence and sub-regions

N1

TCCAGCTCTGAATCTCACTCCCAGGCTCTGCCGAAGGTTCATGGAATCCTTACGCTGAATAATTTCAAC

m1 N2

ATATCGCTTCCGTTTGTCAGCAAATATTTCGGAGTAAAAGCGACCCAACAGAACTAACCCGGCCCTCGA

m2 N3

CGCTAAAATGATAAAAATTACCACTTAATTGGATCGCCCCAATATGCGACAGCTCTCCAGAGATCGAGC

m3

GTCAGTAGGAGGCTTGCCTCCTTTTATTCCTGCGGCAACGAGATAAAAATCTAATTTATAGCTTAGACTA

*Blot*

N4 N5 N6

TTTCGTGTGATTTCGGGACCCATCCACAGGTAGCTGACCGTTCTGATCTCGGCTACAAATGCCCCACAA

m4

CTTAGGTCCTGCCAGCCCGCAATGGATTCATATATTTTCGTATTTAATAAACAGGCTACCGAATTTCGCAT

*Xasta*

TTCCTGCCTACCTTCAAAAGGTCGCTCGATTTTAGAGCTGGCCTCC
