## Supplementary Table 1 for "In situ mutational screening and CRISPR interference define *apterous* cis-regulatory inputs during compartment boundary formation"

| Fly stock | Plasmid used for its generation: | Stock used to inject the plasmids |
|---|---|---|
| *apR2-WT* | MS377 pRMCEentry –attB +yellow +WT in pUC57-Kan | apOR463 landing site |
| *apR2-Δm1* | MS378 pRMCEentry –attB +yellow +m1 in pUC57-Kan | apOR463 landing site |
| *apR2-Δm2* | MS379 pRMCEentry –attB +yellow +m2 in pUC57-Kan | apOR463 landing site |
| *apR2-Δm3* | MS380 pRMCEentry –attB +yellow +m3 in pUC57-Kan | apOR463 landing site |
| *apR2-Δm4* | DB341 pRMCEentry –attB +yellow +m4 in pUC57-Kan | apOR463 landing site |
| *apR2-Δm1m4* | MS381 pRMCEentry –attB +yellow +m1m4 in pUC57-Kan | apOR463 landing site |
| *apR2-Δm1.1m4* | MS383 pRMCEentry –attB +yellow +m1m4 in pUC57-Kan | apOR463 landing site |
| *apR2-Δm1.2m4* | MS354 pRMCEentry –attB +yellow +m1.2m4 in pUC57-Kan | apOR463 landing site |
| *apR2-Δm1.3m4* | MS384 pRMCEentry –attB +yellow +m1.3m4 in pUC57-Kan | apOR463 landing site |
| *apR2-Δm3.1* | MS385 pRMCEentry –attB +yellow +m3.1 in pUC57-Kan | apOR463 landing site |
| *apR2-Δm3.2* | MS386 pRMCEentry –attB +yellow +m3.2 in pUC57-Kan | apOR463 landing site |
| *apR2-Δm3.3* | DB342 pRMCEentry –attB +yellow +m3.3 in pUC57-Kan | apOR463 landing site |
| *apR2-Δm3.4* | MS387 pRMCEentry –attB +yellow +m3.4 in pUC57-Kan | apOR463 landing site |
| *apR2-ΔN1* | MS388 pRMCEentry –attB +yellow +N1 in pUC57-Kan | apOR463 landing site |
| *apR2-ΔN2* | MS389 pRMCEentry –attB +yellow +N2 in pUC57-Kan | apOR463 landing site |
| *apR2-ΔN3* | MS390 pRMCEentry –attB +yellow +N3 in pUC57-Kan | apOR463 landing site |
| *apR2-ΔN4* | MS391 pRMCEentry –attB +yellow +N4 in pUC57-Kan | apOR463 landing site |
| *apR2-ΔN6* | MS392 pRMCEentry –attB +yellow +N5 in pUC57-Kan | apOR463 landing site |
| *apR2-Δm1m2m4* | MS396 pRMCEentry –attB +yellow +m1m2m4 in pUC57-Kan | apOR463 landing site |
| *R2-OR463m3ΔAA* | pRMCEentry –attB +yellow +m3ΔAA in pUC57-Kan | apOR463 landing site |
| *R2-OR463m3+6bp* | pRMCEentry –attB +yellow +m3+6bp in pUC57-Kan | apOR463 landing site |
| *R2-OR463m2Δ6bpm3* | pRMCEentry –attB +yellow +m2Δ6bpm3 in pUC57-Kan | apOR463 landing site |
| *apΔN5* | Isolated as an indel at one of the gRNAs used to stablish the apR2 landing site | |
| m3 substitution library | | |
| *R2-m3.1-A* | pRMCEentry –attB +yellow +m3.1-A in pUC57-Kan | apOR463 landing site |
| *R2-m3.1-B* | pRMCEentry –attB +yellow +m3.1-B in pUC57-Kan | apOR463 landing site |
| *R2-m3.1-C* | pRMCEentry –attB +yellow +m3.1-C in pUC57-Kan | apOR463 landing site |
| *R2-m3.1-D* | pRMCEentry –attB +yellow +m3.1-D in pUC57-Kan | apOR463 landing site |
| *R2-m3.1-E* | pRMCEentry –attB +yellow +m3.1-E in pUC57-Kan | apOR463 landing site |
| *R2-m3.1-F* | pRMCEentry –attB +yellow +m3.1-F in pUC57-Kan | apOR463 landing site |
| *R2-m3.1-G* | pRMCEentry –attB +yellow +m3.1-G in pUC57-Kan | apOR463 landing site |
| *R2-m3.1-H* | pRMCEentry –attB +yellow +m3.1-H in pUC57-Kan | apOR463 landing site |
| *R2-m3.1-I* | pRMCEentry –attB +yellow +m3.1-I in pUC57-Kan | apOR463 landing site |
| *R2-m3.1-J* | pRMCEentry –attB +yellow +m3.1-J in pUC57-Kan | apOR463 landing site |
| *R2-m3.1-K* | pRMCEentry –attB +yellow +m3.1-K in pUC57-Kan | apOR463 landing site |
| *R2-m3.1-L* | pRMCEentry –attB +yellow +m3.1-L in pUC57-Kan | apOR463 landing site |
| *R2-m3.1-M* | pRMCEentry –attB +yellow +m3.1-M in pUC57-Kan | apOR463 landing site |
| *R2-m3.1-N* | pRMCEentry –attB +yellow +m3.1-N in pUC57-Kan | apOR463 landing site |
| *R2-m3.1-O* | pRMCEentry –attB +yellow +m3.1-O in pUC57-Kan | apOR463 landing site |
| *R2-m3.1-P* | pRMCEentry –attB +yellow +m3.1-P in pUC57-Kan | apOR463 landing site |
| *R2-m3.1-Q* | pRMCEentry –attB +yellow +m3.1-Q in pUC57-Kan | apOR463 landing site |
| *R2-m3.1-R* | pRMCEentry –attB +yellow +m3.1-R in pUC57-Kan | apOR463 landing site |
| *R2-m3.1-S* | pRMCEentry –attB +yellow +m3.1-S in pUC57-Kan | apOR463 landing site |
| *R2-m3.1-T* | pRMCEentry –attB +yellow +m3.1-T in pUC57-Kan | apOR463 landing site |
| *R2-m3.1-U* | pRMCEentry –attB +yellow +m3.1-U in pUC57-Kan | apOR463 landing site |
| Other lines | | |
| *apEΔm3-LacZ* | attB-apEΔm3-LacZ | (attP-86Fb)ZH |
| *UAS-dCas9* | pUAStattB-dCas9 | (attP-86Fb)ZH |
| *U6-OR463.gRNAx4* | pCFD5-U6-OR463.gRNAx4 | attp40 (BL25709) |
| *U6-antp.gRNAx3* | pCFD5-U6-antp.gRNAx3 | attp40 (BL25709) |
| *CON2-mKate2* | CON2-mKate2 | (attP-86Fb)ZH |
| *CON5-eGFP-NLS* | CON5-eGFP-NLS | (attP-86Fb)ZH |
