## Supplementary Table 2 for "In situ mutational screening and CRISPR interference define *apterous* cis-regulatory inputs during compartment boundary formation"

| Primer name | Sequence 5'-3' | Plasmid generated with it |
|---|---|---|
| m1.1 (A317) | gttcatggaatccttacgctgaatatcgcttccgtttgtcagcaaatatttcgg | MS383 pRMCEentry –attB +yellow +m1m4 in pUC57-Kan |
| m1.2 (A318) | acgctgaataatttcaacatatgtcagcaaatatttcggagtaaaagc | MS354 pRMCEentry –attB +yellow +m1.2m4 in pUC57-Kan |
| m1.3 (A319) | tttcaacatatcgcttccgttcaaatatttcggagtaaaagcgac | MS384 pRMCEentry –attB +yellow +m1.3m4 in pUC57-Kan |
| m3.1 (A320) | cctccttttattcctgcggcaaccttagactatttcgtgtgatttcgg | MS385 pRMCEentry –attB +yellow +m3.1 in pUC57-Kan |
| m3.2 (A321) | ctccttttattcctgcggcaacaatctaatttatagcttagactatttcg | MS386 pRMCEentry –attB +yellow +m3.2 in pUC57-Kan |
| m3.3 (A322) | tttattcctgcggcaacgagataaaatagcttagactatttcgtgtg | DB342 pRMCEentry –attB +yellow +m3.3 in pUC57-Kan |
| m3.4 (A323) | gcaacgagataaaaatctaatttcttagactatttcgtgtgatttcgg | MS387 pRMCEentry –attB +yellow +m3.4 in pUC57-Kan |
| m1 Mutagenic Primer (A267) | gttcatggaatccttacgctgaatacaaatatttcggagtaaaagcgac | MS378 pRMCEentry –attB +yellow +m1 in pUC57-Kan & MS381 (m1m4) |
| m2 Mutagenic Primer (A268) | aagcgacccaacagaactaacccgccccaatatgcgacagctc | MS379 pRMCEentry –attB +yellow +m2 in pUC57-Kan |
| m3 Mutagenic Primer (A269) | gcttgcctccttttattcctcttagactatttcgtgtgatttcgg | MS380 pRMCEentry –attB +yellow +m3 in pUC57-Kan |
| m4 Mutagenic Primer (A270) | cgcaatggattcatatattttcgttaccttcaaaaggtcgctcg | DB341 pRMCEentry –attB +yellow +m4 in pUC57-Kan & MS381 (m1m4) |
| N1 mutagenetic primer (P43) | tgaatctcactcccatttcaacatatcgc | MS388 pRMCEentry –attB +yellow +N1 in pUC57-Kan |
| N2 mutagenetic primer (P44) | cttccgtttgtcaggccctcgacgctaaaatg | MS389 pRMCEentry –attB +yellow +N2 in pUC57-Kan |
| N3 mutagenetic primer (P45) | acttaattggatcggcggcaacgagataaaaatc | MS390 pRMCEentry –attB +yellow +N3 in pUC57-Kan |
| N4 mutagenetic primer (P46) | aatctaatttatagcatccacaggtagc | MS391 pRMCEentry –attB +yellow +N4 in pUC57-Kan |
| N6 mutagenetic primer (P47) | acaggtagctgaccatttaataaacagg | MS392 pRMCEentry –attB +yellow +N5 in pUC57-Kan |
