## Supplementary Figures for "In situ mutational screening and CRISPR interference define *apterous* cis-regulatory inputs during compartment boundary formation"

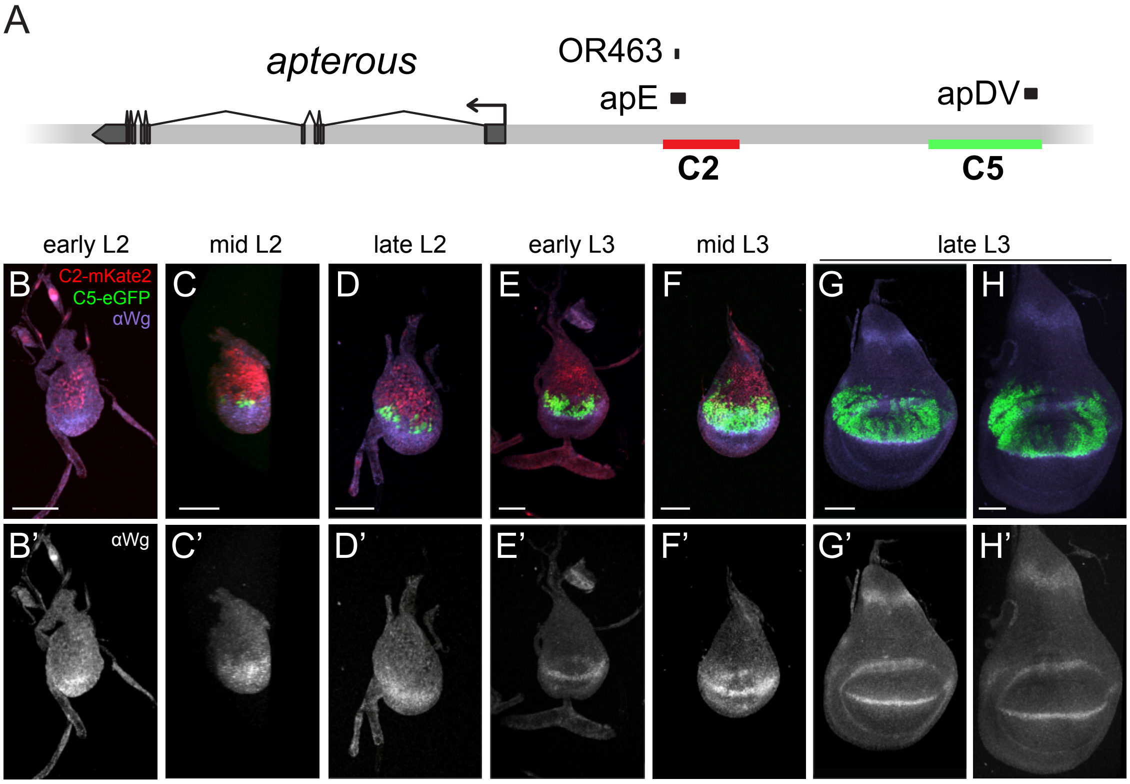


**Figure 1 – figure supplement 1. C2 and C5 activity throughout larval development. A.** illustration of *ap* locus. *ap* coding region is highlighted in dark grey, C2 in red and C5 in green. Within them, apE, OR463 and apDV are labelled in black. **B-H.** Wing discs of C2-mKate2 (red) and C5-eGFP (green) animals dissected at the indicated stages and immunostained for anti-Wg (blue). In early stages, only C2 is active in the central part of the tissue (B). C2 activity peaks in mid L2 (C) to later decrease and eventually disappear (G-H). C5 activity starts in few cells at the DV boundary (C) and later expands to all the dorsal pouch and part of the dorsal hinge (D-H). **B'-H'.** anti-Wg signal of the panels B-H. Scale bars: B-F: 20µm G-H: 50µm.

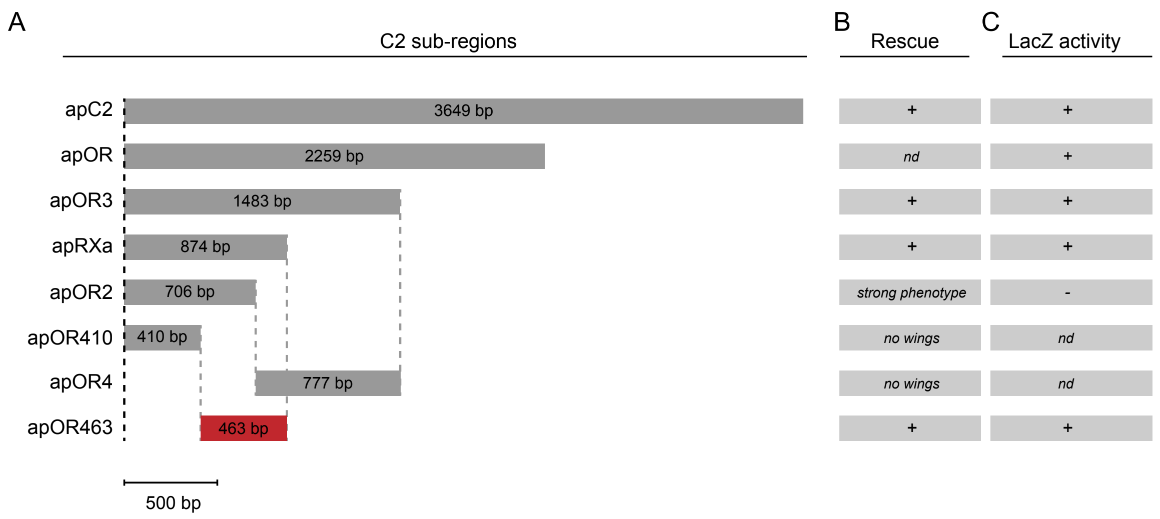


**Figure 1 – figure supplement 2. Identification of the minimal apE enhancer within apC2. A.** Eight DNA fragments from the apC2 region are shown. Their names are given on the left of the panel and their length in bp is indicated. These fragments were tested with two in vivo assays. (1) The fragments were introduced into reentry plasmid DB59 where they are combined with apC5 (contains apDV). These plasmids were brought back into landing site $ap^{attP\Delta Enh}$ (Bieli et al 2015). Stocks were established and their rescue potential was scored in hemizygous flies (over $ap^{DG3}$). (2) The fragments were cloned into a LacZ reporter plasmid and inserted into the zh-86Fb landing site. Their LacZ activity was assayed by standard procedures. **B.** The rescue activity of each allele is shown. +: normal wings are formed. nd = not done. **C.** The LacZ activity of each reporter construct is shown. +: normal apterous expression pattern in the wing imaginal disc. -: no LacZ activity. nd = not done. Based on these experiments, fragment OR463 was chosen as the minimal apE fragment.

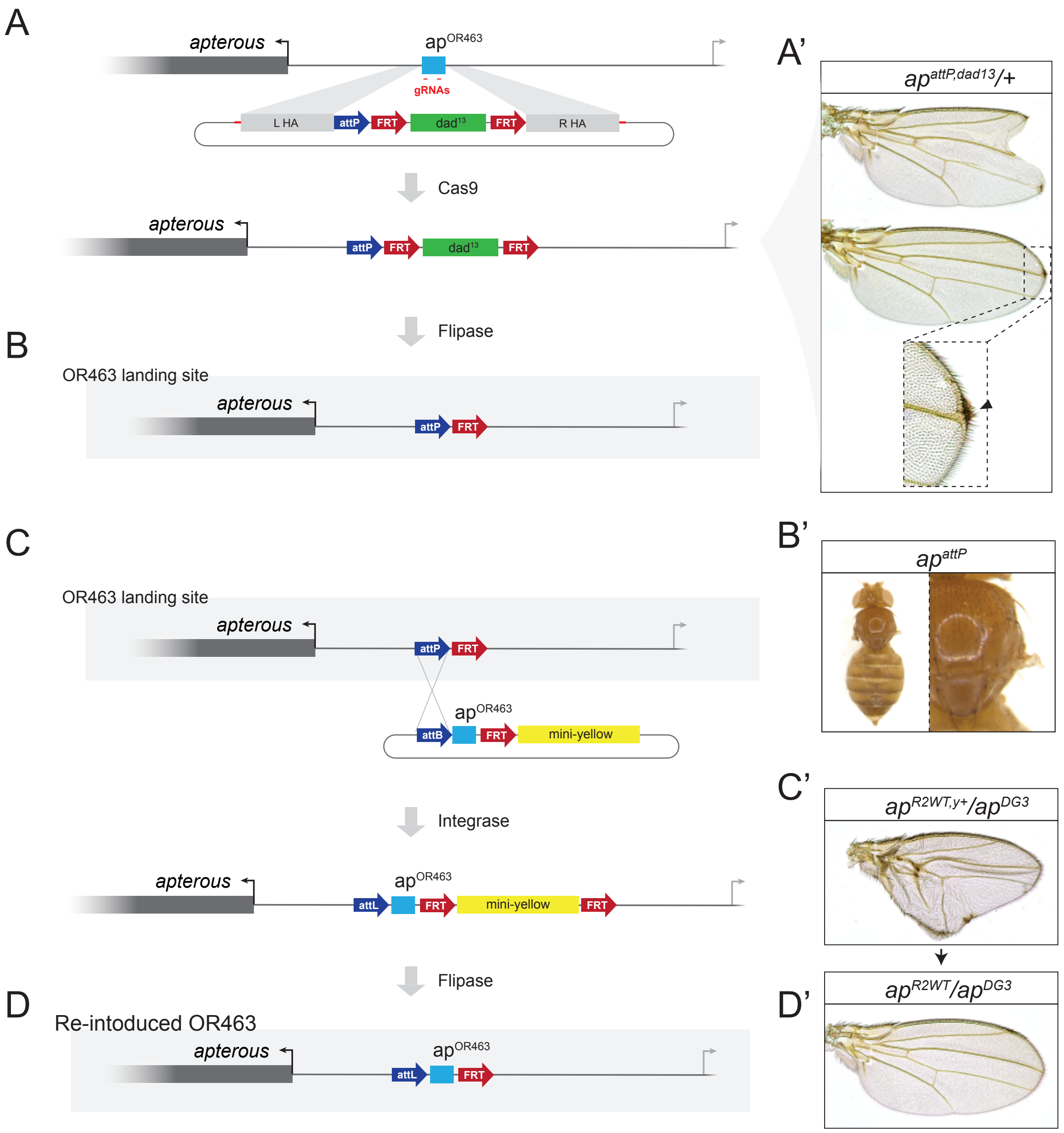


**Figure 1 – figure supplement 3. Generation and validation of the OR463 landing site. A.** Schematic representation of CRISPR- mediated targeting of OR463. Embryos were injected with plasmids encoding for two gRNAs (red) laying within the OR463 fragment (blue box) to direct Cas9 to the locus. A donor plasmid was co-injected to serve as template during HDR. The donor plasmid contained: 1) the *attP* site (dark blue), 2) Dad13 minimal enhancer (517bp in green), flanked by FRT sites (red) in identical orientation, 3) homology arms (each 536bp long) to trigger integration via HDR, 4) flanking gRNA targets to linearize the donor plasmid inside the embryos. **A'.** Dominant wing phenotype of animals in which dad13 was integrated in *ap* locus. These animals presented variable phenotypes despite all having the same insertion. Phenotypes ranged from almost wild-type wings with small border defects in the distal tip (Lower wing, with detail of the tip) to a partial phenocopy of the classical Xasta indentations (upper wing). **B.** Dad13 was then excised by the action of Flp. **B'.** Homozygous wing phenotype of flies in which OR463 has been substituted by the landing site. **C.** Scheme of the re-integration of wild-type OR463 into the landing site. Re-integration constructs contained: 1) an *attB* site, 2) the OR463 sequence, 3) an FRT oriented as that in the landing site, 4) a mini-yellow marker. After insertion via φ C31 integrase, $y^+$ transformants were used to establish stocks. In a second step, the $y^+$ marker was deleted by Flp tratment. **C'.** Hemizygous wing phenotype of flies containing the reintegrated WT OR463 sequence before removal of the mini-yellow marker. These wings often presented aberrant phenotype, which included mirror-image A wing duplications. **D.** Final arrangement of the locus after Flp-mediated excision of mini-yellow. **D'.** Wing phenotype when wild-type OR463 sequence was re-inserted in the OR463 landing site after mini-yellow excision (in hemizygosis).

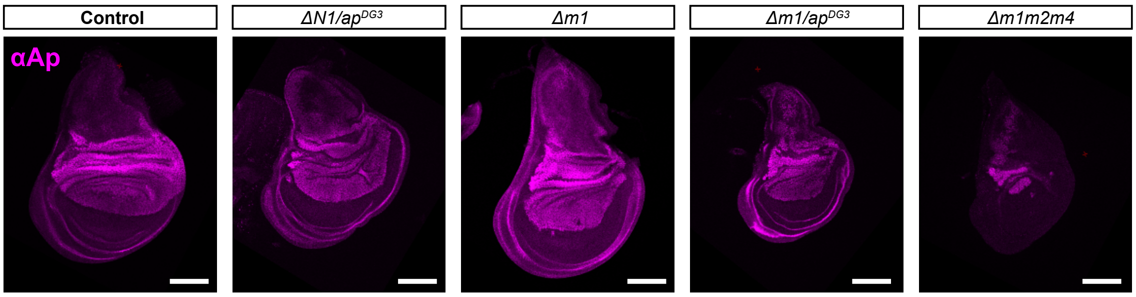


**Figure 2 – figure supplement 1. Ap localization in various genotypes.** Anti-Ap immunostaining in different OR463 mutants. Notice the deformation of the DV boundary in the P compartment in *ΔN1/ ap*$^{DG3}$, *Δm1* and *Δm1/ap*$^{DG3}$. In *Δm1m2m4* (note that m3 is still present in this genotype), the Ap expression pattern is severely reduced, apparently missing from the P compartment**.** Scalebars: 100μm

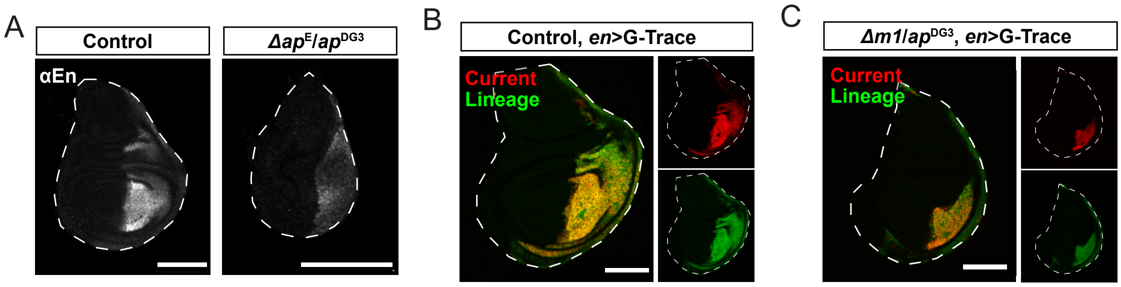


**Figure 2 – figure supplement 2. Loss of *en* expression cannot explain the mirror-image duplications of apE mutants. A.** Anti-En staining in wild-type and *Δap*$^{E}$ mutants in hemizygosis (*Δap*$^{E}$/ *ap*$^{DG3}$). While the wing disc size of *Δap*$^{E}$ mutants is affected, En localization is still constrained to the P side and there is not dramatic change in its pattern**. B.** Control and **C.** *Δm1* hemizygous wing discs in which the current expression of *en* (Red) and its history of expression (Green) are overlayed. No big differences are found between the two signals apart from small clones of current expression present in both control and mutant. *Δm1* hemizygous wing discs present a severe deformation of AP boundary and the P size is severely affected. Scalebars: 100μm

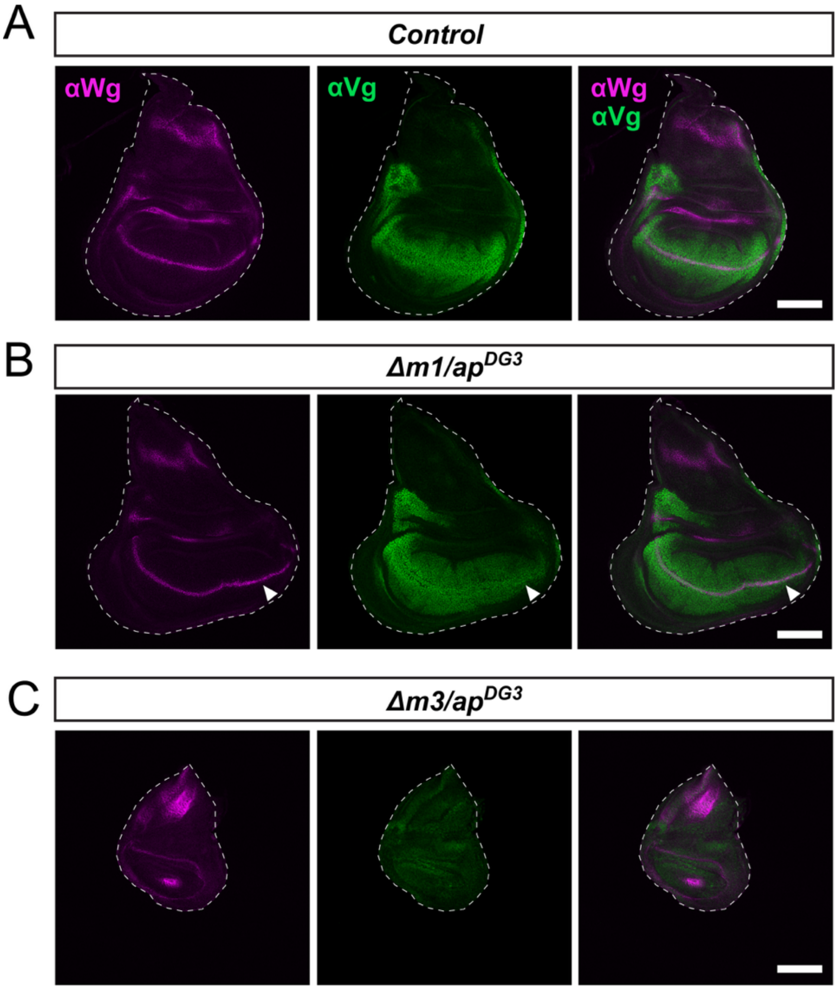


**Figure 2 – figure supplement 3: Vestigial expression in OR463 mutants.** In magenta, anti-Wg immunostaining, in green anti-Vg. **A.** Control (R2-WT/apDG3) L3 wing disc. Wg is detected in its stereotypic stripe along the wing pouch and is also detected in the hinge and notum. Vg is detected in the wing pouch and in the laterals of the proximal hinge. **B.** Hemizygous *Δm1* mutant (*R2-Δm1/apDG3*) L3 wing disc. Wg stripe is extended in the anterior compartment (arrow) coinciding with the outgrowth. Vg localizes throughout the outgrowth in a pattern highly reminiscent of the pouch staining, with its highest levels in along the DV boundary. **C.** Hemizygous Δm3 mutant (*R2-Δm3/apDG3*) L3 wing disc. Wg stripe is missing and only the outer Wg ring of the hinge is detected, the inner ring being reduce to a group of cells. Vg is not detected at levels comparable to the WT discs.

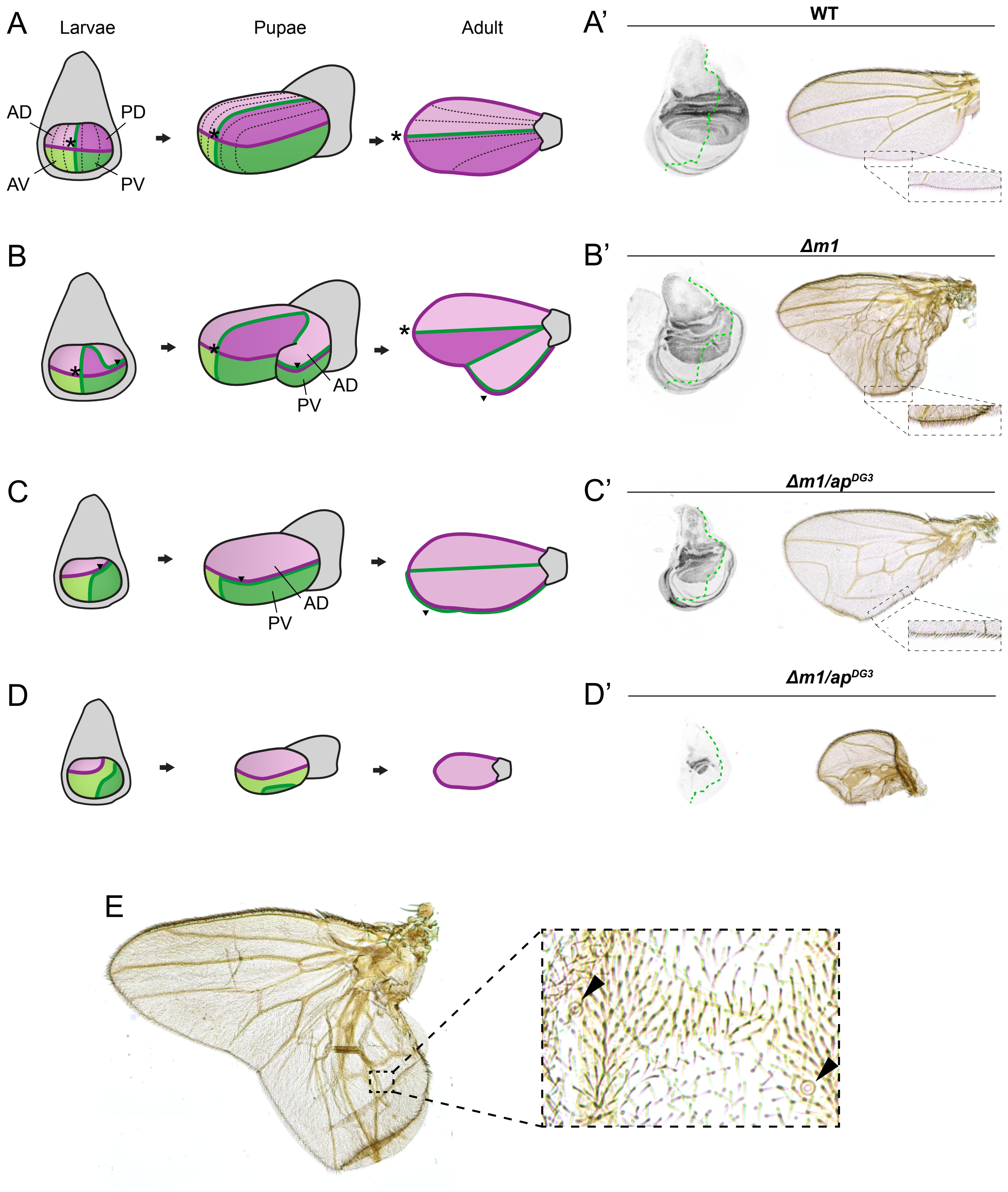


**Figure 2 – figure supplement 4. Model for the development of mirror image duplications in ap OR463 mutants. A.** Model of Wild type wing development across larval, pupal and adult stages. The AP boundary (green) intersect with the DV boundary (purple) in only one spot (asterisk) during larval stages. In pupal stages, the wing pouch everts along the DV boundary, that become the wing margin. As wing veins are patterned by the AP border, their location is relatively parallel to it. **A'.** Example of wild-type wing disc and wing extracted from figure 1 and 2. The morphology of the posterior border is highlighted. It consists of a single row of fine bristles and looks distinctively different from the double-row bristles near of the tip of the. Wing or the triple-row bristles at the anterior margin. B. Wing disc model in which the PD size is affected but still occupies part of the pouch. In these cases, the AP border intersect with the DV boundary in two points which are far enough to specify two independent pouches. The "secondary" pouch does not present a real intersection, but both AP and DV boundaries coincide for a stretch after meeting. During pupal stages, after wing disc eversion, the outgrowth will present a dorsal side with A identity and a ventral side with P identity, resulting in venation defects. These veins, moreover, will be patterned parallel to the margin, as Hh and Dpp will be expressed along the DV boundary. **B'.** Example of a *Δm1* wing disc and wing extracted from figure 1 and 2 that might be following this model. Note the presence of tip-margin bristles in the posterior outgrowth **C.** Wing disc in which PD is almost missing. In these cases, the whole P compartment will behave as does the outgrowth in panel B. **C'.** Example of a *Δm1*/$ap^{DG3}$ wing disc and wing extracted from figure 1 and 2 that might be following this case. In this case, A-specific dorsal triple-row bristles can be detected at the posterior margin. **D.** Model of wing development in which the DV is totally missing. In these cases, DV and AP boundaries are no longer crossing each other. If in close proximity, as it is the case here, a small wing pouch will still be specified. During wing disc eversion, the DV boundary will only present A identity, P compartment being reduced to one of the two sides of the wing depending on the P size, it might form a small "bag-like" structure. **D'** Example of a *Δm1*/DG3 wing disc and wing extracted from figure 1 and 2 that might be following this model. Note that the margin of this wing is only composed of anterior and tip-specific margin bristles **E.** *Δm1* homozygous mutant wing and detail of the campaniform sensillae in the P wing.

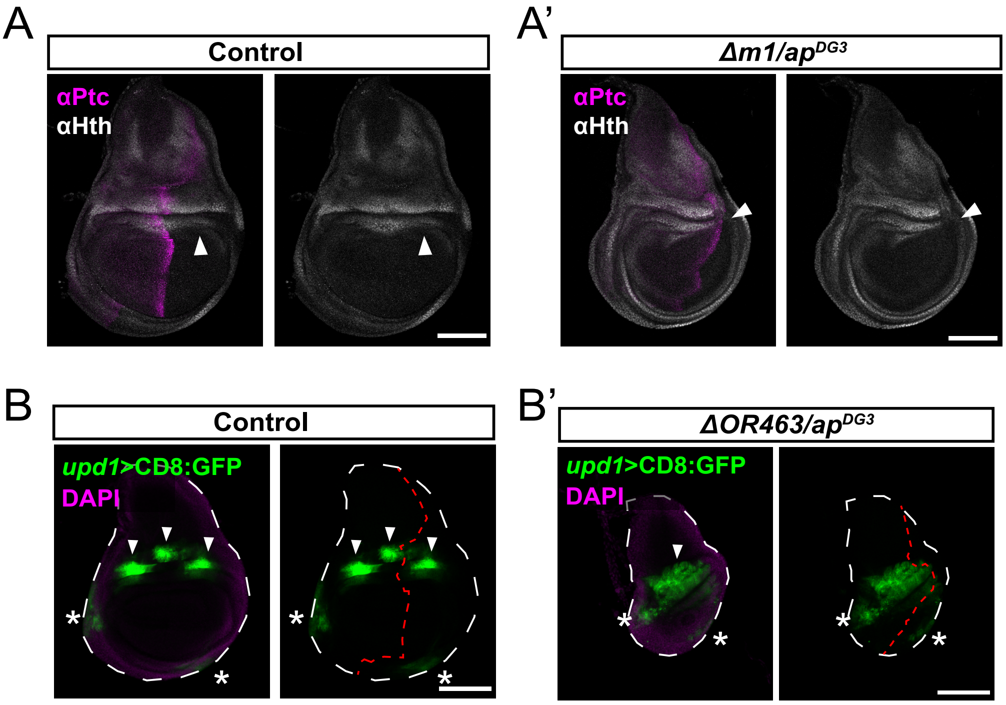


**Figure 2 – figure supplement 5. Posterior hinge and Upd expression patterns are affected in *ap* mutants.** **A**. Locali-zation of Ptc (Magenta) and Hth (Gray) in control wing discs. Hth is expressed along the hinge. (Posterior dorsal hinge is indicated with an arrowhead) **A'.** Localization of Ptc and Hth in *Δm1/ap$^{DG3}$* mutants. In these cases, P hinge is depleted (arrowhead) and the Ptc stripe deviated. **B.** *UAS*-CD8:GFP driven by *upd-Gal4* in wild-type wing discs. In the control discs, GFP signal could be detected in three main spots in the dorsal hinge (arrowheads) and two weak areas of expression in the ventral body wall (asterisks). **B'.** *UAS*-CD8:GFP driven by *upd-Gal4* in *ΔOR463/ ap$^{DG3}$* hemizygous background. Notice that the P source of Upd in the hinge is totally lost in this condition. Scale bars: 100µm

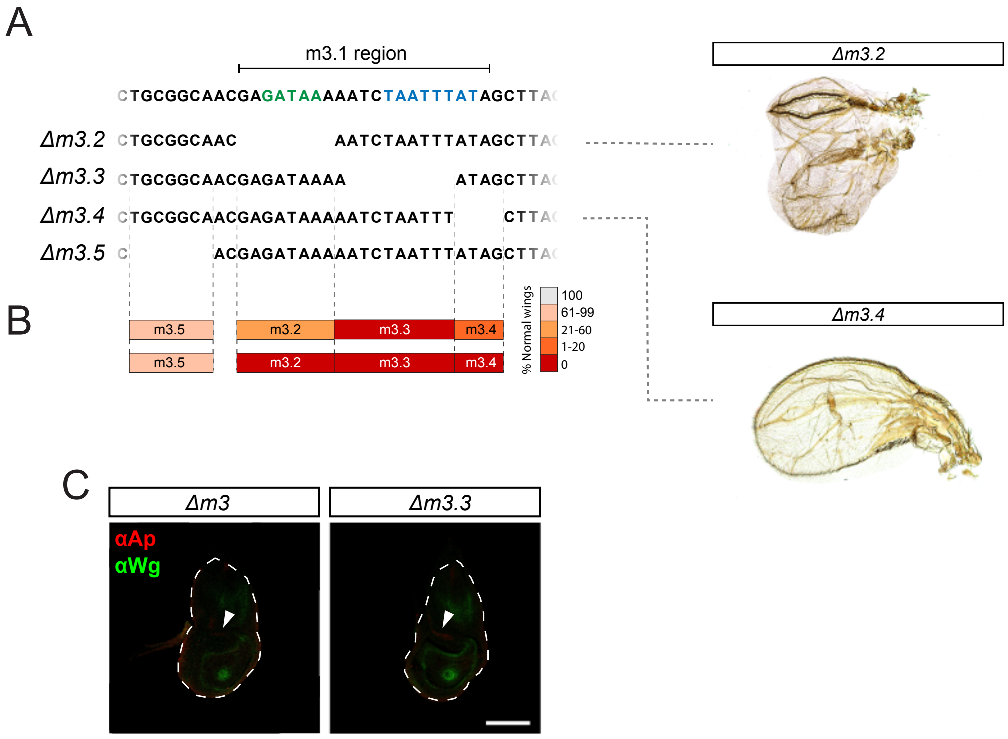


**Figure 5 – figure supplement 1. Deletion analysis within m3 region. A.** Schematic representation of the different deletions and representative phenotypes. Examples of the loss of the GATA binding site and the most distal m3 region are shown. Note that *Δm3*.3 flies do not develop wing structures. **B.** Scoring of the penetrance of mutant phenotypes in the different deletions. **C.** anti-Ap and anti-Wg immunostaining in both *Δm3* deletion and *Δm3.3*. In both cases Ap localization is only residual within the anterior hinge.

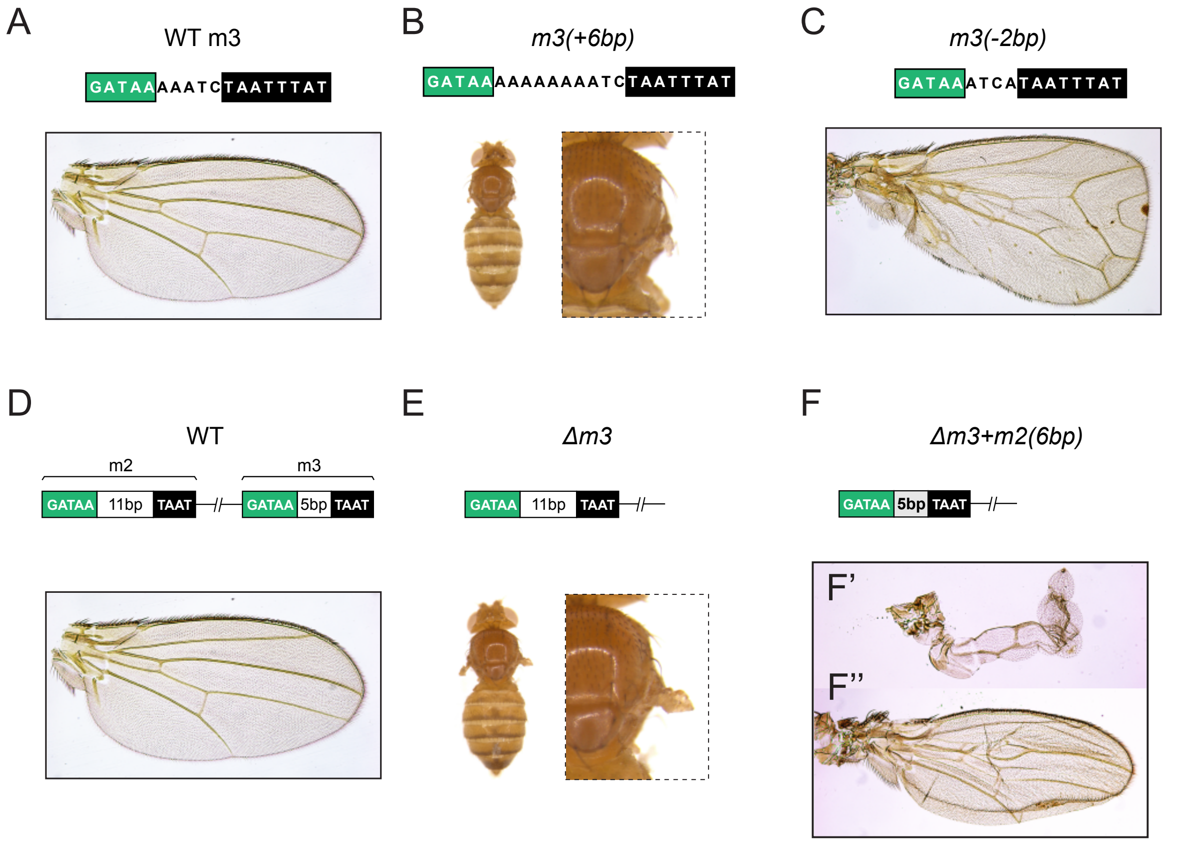


**Figure 5 – figure supplement 2. Correct spacing between GATA and HOX binding sites is essential. A.** Adult wing phenotype of control flies. **B.** Loss of wing produced upon extension of the spacing between the GATA and HOX binding sites by 6pb Notice that this genotype never resulted in normal wings (See scoring of Figure 5D). Wings were either completely missing (35%) or showed sever phenotype. **C.** Example wing phenotype resulting from the contraction of the linker between the GATA and HOX binding sites. In this case, observed phenotypes were clearly weaker than for m3+6bp. Apart from normal wings (30%), most of the wings had an outgrowth from the posterior compartment or looked as depicted. **D.** Adult wing phenotype of control flies (Same specimen as in A). **E.** Loss of wing produced upon *Δm3* deletion (same as Figure 1G). **F.** Example of the wing phenotypes obtained upon contraction of the m2's GATA-HOX linking sequence in the absence of m3. Note that the majority (68%) of wings were missing for this genotype. Among the rest, most wings looked as shown in F'. A few cases as shown in F'' were also present.

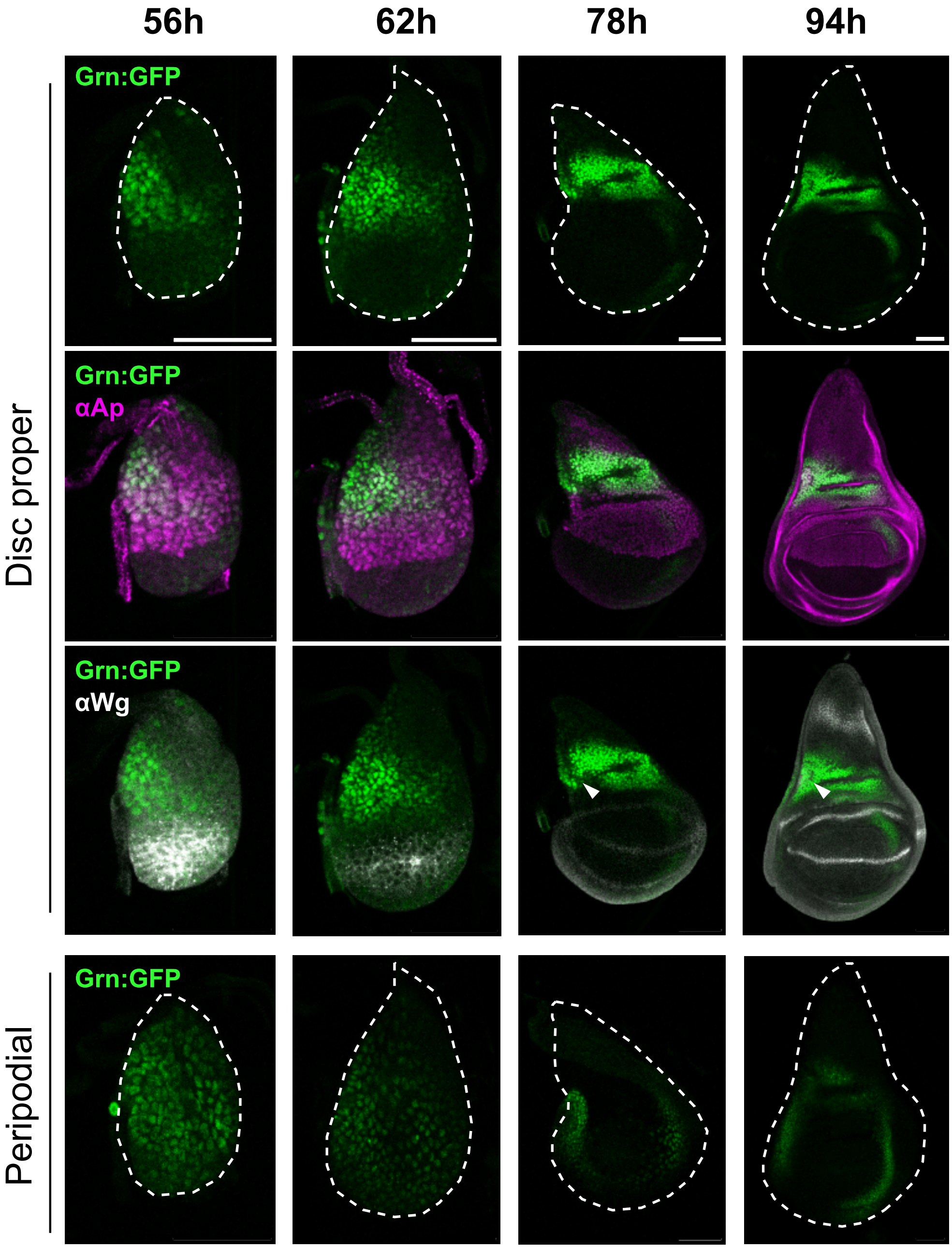


**Figure 6 – figure supplement 1. Pattern of Grn:GFP localization during wing development.** *grn:*GFP wing disc stained with anti-Ap and anti-Wg. (56h) At this stage Grn:GFP localizes to the presumptive hinge and seems to be highly down- regulated in the future notum and incipient pouch (marked by Wg). Grn:GFP was detected at higher levels in the anterior compartment. It was also detected broadly in the peripodial membrane. (62h) Similar Grn:GFP localization to that of 56h. (78h) Grn:GFP localization is higher in the proximal hinge, but persist at low levels in some lateral areas. Peripodial expression is higher in the lateral sides. (94h) Grn:GFP localizes sharply in the proximal hinge, totally excluded from the notum. Proximal hinge levels are very low. Only the cubic cells of the peripodial membrane seem to express *grn*. Arrowheads indicate the intersection between trachea and the disc proper. Scalebars: 50μm

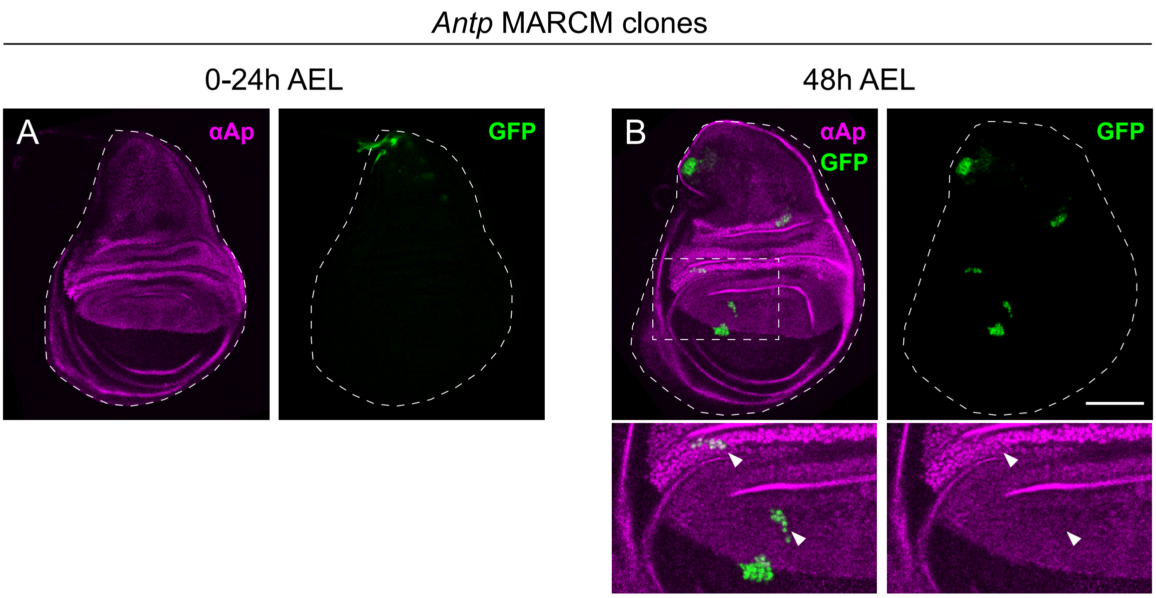


**Figure 7 – figure supplement 1. *Antp* is required for clone survival in early stages and does not affect *ap* expression at later stages. A.** Example of wing disc in which *Antp* MARCM clones were generated during embryonic stages (0-24h) (See materials and methods for full genotype). In this case, many clones could be retrieved in other tissues (data not shown), but never in the disc proper of the wing disc. Clones, marked by the presence of GFP, could only be retrieved in the peripodial membrane (detail of the peripodial membrane in the right panel). In magenta, anti-Ap immunostaining reveals normal Ap pattern (left panel). **B.** Example of a wing disc of the same genotype as in A in which the clones were generated at 48h in development. In this case, small *Antp* clones (GFP positive) could be detected in the disc proper. Anti-Ap immunostaining revealed no change in the levels inside the clone (arrowheads).
